# PLM-interact: extending protein language models to predict protein-protein interactions

**DOI:** 10.1101/2024.11.05.622169

**Authors:** Dan Liu, Francesca Young, Kieran D. Lamb, Adalberto Claudio Quiros, Alexandrina Pancheva, Crispin Miller, Craig Macdonald, David L Robertson, Ke Yuan

**Affiliations:** MRC-University of Glasgow Centre for Virus Research, Glasgow, United Kingdom; School of Cancer Sciences, University of Glasgow, Glasgow, United Kingdom; Cancer Research UK Scotland Institute, Glasgow, United Kingdom; School of Computing Science, University of Glasgow, Glasgow, United Kingdom

## Abstract

Computational prediction of protein structure from amino acid sequences alone has been achieved with unprecedented accuracy, yet the prediction of protein-protein interactions (PPIs) remains an outstanding challenge. Here we assess the ability of protein language models (PLMs), routinely applied to protein folding, to be retrained for PPI prediction. Existing PPI prediction models that exploit PLMs use a pre-trained PLM feature set, ignoring that the proteins are physically interacting. Our novel method, PLM-interact, goes beyond a single protein, jointly encoding protein pairs to learn their relationships, analogous to the next-sentence prediction task from natural language processing. This approach provides a significant improvement in performance: Trained on human-human PPIs, PLM-interact predicts mouse, fly, worm, *E. coli* and yeast PPIs, with 16-28% improvements in AUPR compared with state-of-the-art PPI models. Additionally, it can detect changes that disrupt or cause PPIs and be applied to virus-host PPI prediction. Our work demonstrates that large language models can be extended to learn the intricate relationships among biomolecules from their sequences alone.

## Introduction

Proteins are the main structural components of cells and mediate biological processes by interacting with other proteins^1^. Disruption of these protein-protein interactions (PPIs), e.g., mediated by mutations can underlie human disease^2^. In virology PPIs are particularly important as viruses depend entirely on the host cell for replication, achieved mainly through specific interactions with host proteins. Understanding PPI mechanisms offers the potential for developing novel therapy strategies for both human disease and pathogen infections^3^. Unfortunately, experimentally identifying PPIs is both costly and time-consuming, such that interaction datasets remain sparse with only a few species having comprehensive coverage^4,5^.

Computational algorithms offer an efficient alternative to the prediction of PPIs at scale. Existing prediction approaches mainly leverage protein properties such as protein structures, sequence composition and evolutionary information^6,7,8,9^. Applying these features to pairs of proteins, classifiers have been trained using classical machine learning^10^ and deep learning approaches^11^. Recently, protein language models (PLMs) trained on large public protein sequence databases have been used for encoding sequence composition, evolutionary and structural features^12–14^, becoming the method of choice for representing proteins in state-of-the-art PPI predictors. A typical PPI prediction architecture uses a pre-trained PLM to represent each protein in a pair separately, then a classification head is trained for a binary task that discriminates interacting pairs from non-interacting pairs^13,15^ (**Figure 1a**). Despite this use of PLMs, identifying positive PPIs remains challenging.

**Figure 1.**
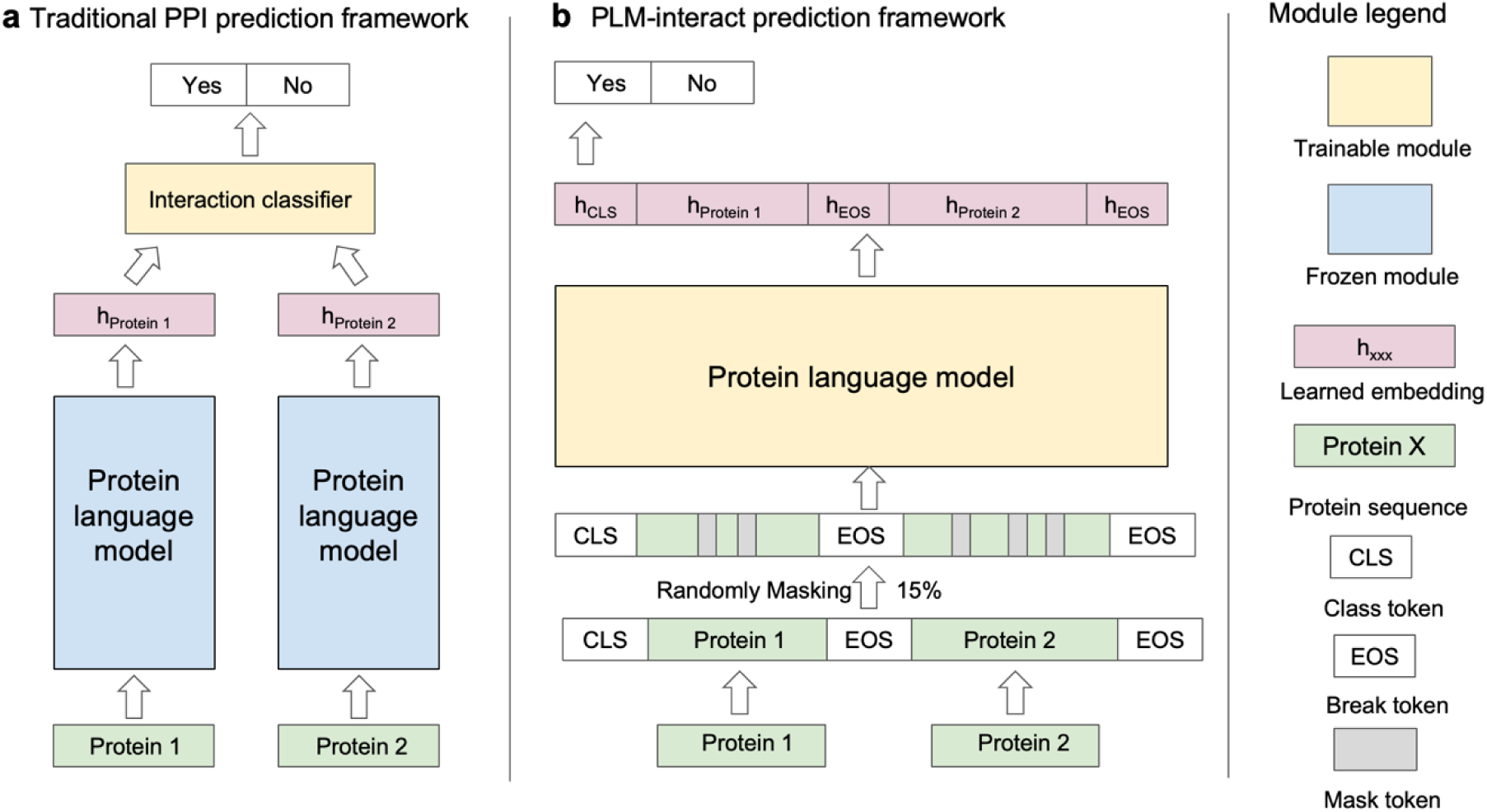
A comparison of PLM-interact to the typical existing PPI prediction framework. **a.** PPI prediction models that use pre-trained protein language models to extract single protein embeddings. Then an interaction classifier is trained use these single protein embeddings. **b.** PLM-interact uses a protein language model with a longer context to handle a pair of protein sequences directly. Both the mask language modelling task and a binary classification task predicting interaction status are used to train the model. (see **Supplementary Figure 1**).

The main issue is PLMs are primarily trained using single protein sequences, i.e., while they learn to identify contact points within a single protein^16^, they are not ‘aware’ of interaction partners. In a conventional PLM-based PPI predictor architecture, a classification head is used to extrapolate the signals of inter-protein interactions by grouping common patterns of intra-protein contacts in interacting and non-interacting pairs, respectively (**Figure 1a**). This strategy relies on the classification head being generalisable. Unfortunately, with a feedforward neural network being the dominant option, these classifiers often don’t generalise well.

To address the lack of inter-protein context in training, we propose a novel PPI prediction model, PLM-interact, that directly models PPIs by extending and fine-tuning a pre-trained PLM, ESM-2^17^. PLM-interact (trained on human PPI data) achieves a significant improvement compared to other predictors when applied to mouse, fly, worm, yeast and *E. coli* datasets. We demonstrate that PLM-interact is capable at identifying mutations that cause and disrupt interactions. We also show PLM-interact’s performance in a virus-human PPI prediction task.

### PLM-interact

To directly model PPIs, two extensions to ESM-2 are introduced (**Figure 1b**): (1) longer permissible sequence lengths in paired masked-language training to accommodate amino acid residues from both proteins; (2) implementation of the “next sentence” prediction task^18^ to fine-tune ESM-2 where the model is trained with a binary label indicating whether the protein pair is interacting or not (see Methods for more details). Our training task is, thus, a mixture of the next sentence prediction and the mask language modelling tasks. This architecture allows amino acids in a protein to be linked by amino acids from a different protein through transformer’s attention mechanism.

The training of PLM-interact begins with the pre-trained large language model ESM-2. We fine-tune it for PPIs, by showing it pairs of known interacting and non-interacting proteins. In contrast to similar training strategies in machine learning^18^, we find the next sentence prediction and mask language modelling objectives need to be balanced. We therefore conducted comprehensive benchmarking for different weighting options, before selecting a 1:10 ratio between classification loss and mask loss, combined with initialisation using the ESM-2 (with 650M parameters), as this achieved the best performance (see Methods and **Supplementary Figure 1).**

### PLM-interact improves prediction performance

To examine the performance of PLM-interact, we benchmark the model against five PPI prediction approaches: TT3D^13^, Topsy-Turvy^19^, D-SCRIPT^15^, PIPR^6^ and DeepPPI^20^. We use a multi-species dataset created by Sledzieski et al.^15^. Each model is trained on human protein interaction data and tested on five other species. The human training dataset in this multi-species dataset includes 421,792 protein pairs (38,344 positive interaction pairs and 383,448 negative pairs), human validation includes 52,725 protein pairs (4,794 positive interaction pairs and 47,931 negative pairs) and the mouse, worm, fly and yeast test datasets each includes 55,000 pairs (5,000 positives interaction pairs and 50,000 negative pairs), except for the *E. coli* test dataset, which includes 22,000 pairs (2,000 positive interaction pairs and 20,000 negative pairs). The positive PPIs in these datasets are experimentally-derived physical interactions, while the negative pairs are randomly paired proteins not reported to interact.

PLM-interact achieves the highest AUPR (area under the precision-recall curve)^23^ with the next best performer, TT3D^13^, although similar performance to Topsy-Turvy^19^ (**Figure 2b**). Tested on mouse, fly and worm test species datasets, PLM-interact has an improvement of 16%, 21% and 20% AUPR compared to TT3D^13^, respectively. The predictions for yeast and *E. coli* PPIs are more challenging because they are more evolutionarily divergent from the human proteins used for training than the other species (see **Figure 2b**): Our model achieved an AUPR of 0.706 on yeast, a 28% improvement over TT3D’s AUPR of 0.553 and a 19% improvement on *E. coli* with an AUPR of 0.722.

**Figure 2.**
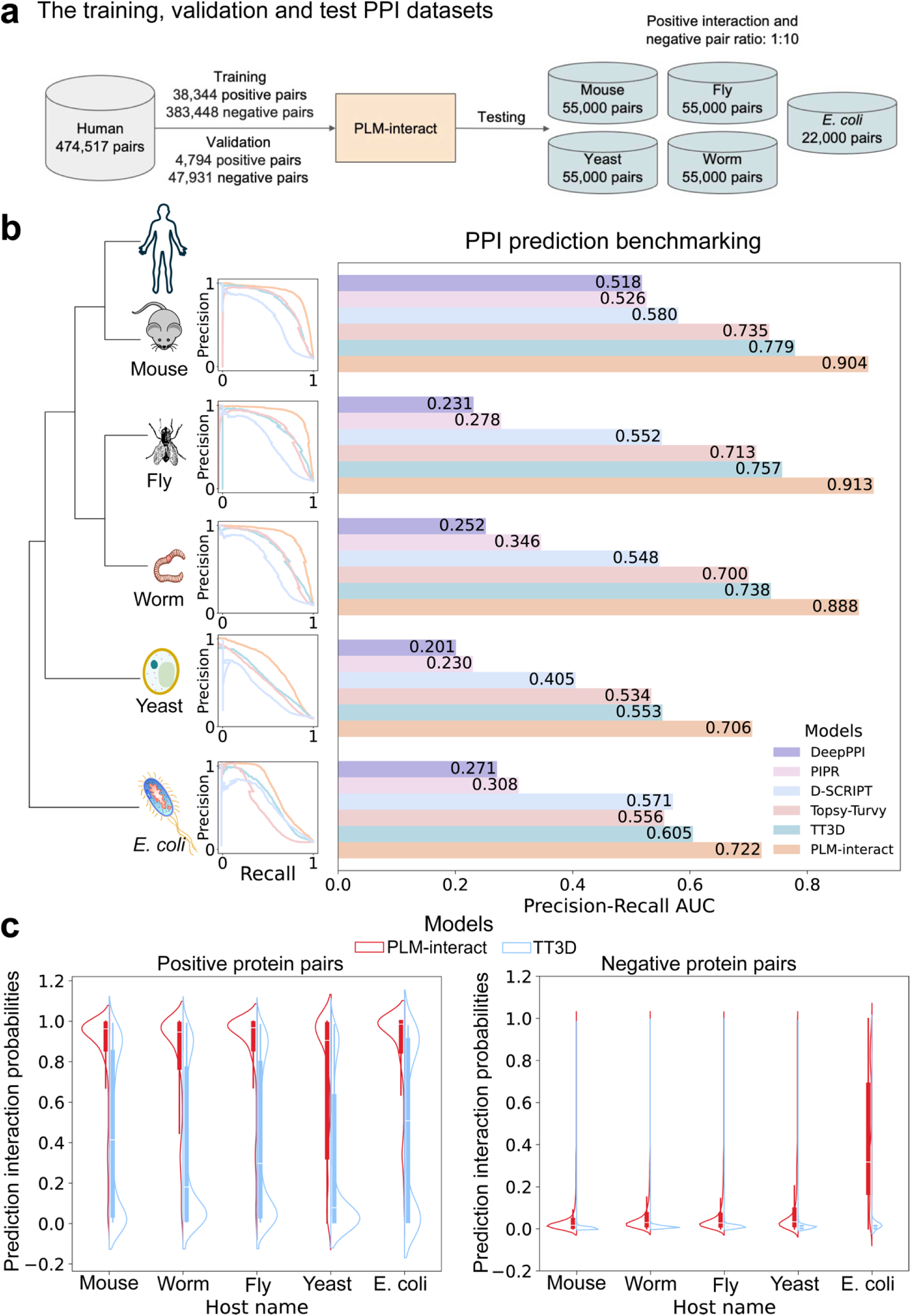
PLM-interact achieves the highest PPI prediction performance. The benchmarking results of PLM-interact with state-of-the-art PPI prediction models. **a.** The data size of training, validation and test PPIs. **b.** The taxonomic tree of the training and test species, precision-recall curves of each test species and a bar plot of AUPR values on PPI prediction benchmarking. **c**. Violin plots of predicted interaction probabilities of PLM-interact and TT3D on positive and negative pairs, respectively. (see **Supplementary Figure 2** for more information).

Importantly, the improvement in PLM-interact is due to its ability to correctly identify positive PPIs: Comparing the prediction interaction probability of PLM-interact with the second-best predictor TT3D, PLM-interact consistently assigned higher probabilities of interaction to true positive PPIs. TT3D, in contrast, despite using a broader feature set, produces a bimodal distribution for interaction probabilities in all held-out species (more details see **Supplementary Figure 2**).

Next, we showcase five positive PPI instances, one for each test species, for which our model produces a correct prediction, but TT3D produces an incorrect prediction (**Figure 3**). These PPIs are necessary for essential biology processes including ubiquinone biosynthesis, RNA polymerisation, ATP catalysis, transcriptional activation and protein transportation. We use Chai-1^21^ and AlphaFold3^22^ to predict and visualise these interacting protein structures (**Figure 3** and **Supplementary Figure 3**). Notably, both Chai-1 and AlphaFold3 have only 1 out 5 structures with close to high-confidence prediction (ipTM close to 0.8). Both methods give failed prediction scores (ipTM <0.6) for 4 out 5 structures. PLM-interact gives correct predictions with high confidence in all cases.

**Figure 3.**
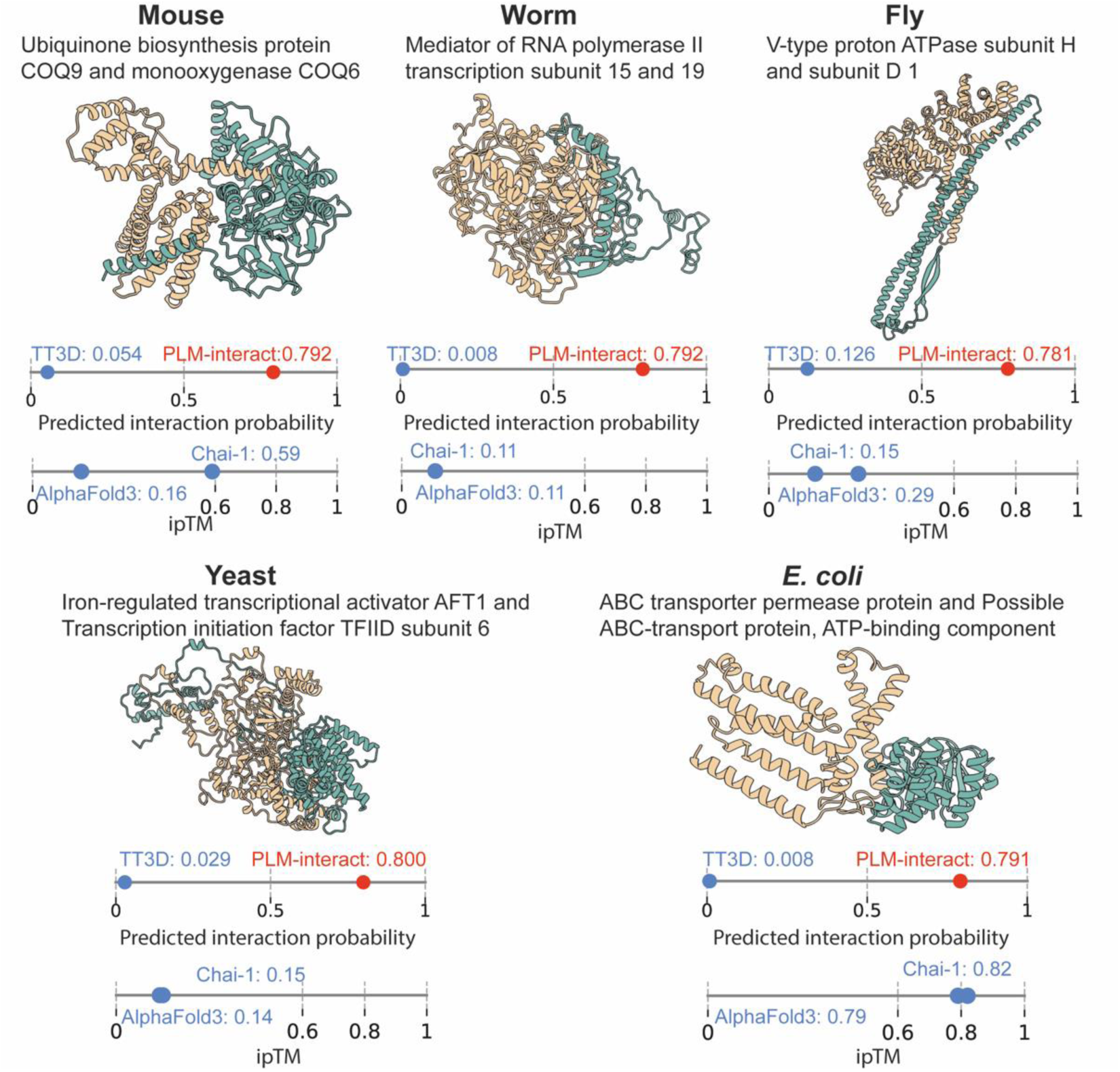
PPI example for each species that was predicted correctly by PLM-interact but not by TT3D. Protein-protein structures are predicted by Chai-1^21^ and visualised with ChimeraX^23^. Both models’ prediction interaction probabilities range between 0 and 1. A predicted interaction probability >0.5, is predicted as a positive PPI, while <0.5 is a negative pair. Interacting proteins are shown from left (yellow) to right (green), respectively, for **mouse**: Q8K1Z0 (Ubiquinone biosynthesis protein COQ9, mitochondrial) and Q8R1S0 (Ubiquinone biosynthesis monooxygenase COQ6, mitochondrial); **Worm**: Q21955 (Mediator of RNA polymerase II transcription subunit 15) and Q9N4F2 (Mediator of RNA polymerase II transcription subunit 19); **Fly**: Q9V3J1 (V-type proton ATPase subunit H) and Q9V7D2 (V-type proton ATPase subunit D 1); **Yeast**: P22149 (Iron-regulated transcriptional activator AFT1) and P53040 (Transcription initiation factor TFIID subunit 6); and ***E. coli***: A0A454A7G5 (ABC transporter permease protein) and A0A454A7H5 (Possible ABC-transport protein, ATP-binding component). See **Supplementary Figure 3** for AlphaFold3^22^ predicted structures. The ipTMs for both Chai-1 and AlphaFold3 are shown for each structure, where ipTM <0.6 indicates failed predictions and ipTM >0.8 indicates high confidence predictions.

### PLM-interact can identify the impact of mutations on interactions

Here, we demonstrate examples of PLM-interact correctly predicting the consequences of mutations on PPIs. Canonical protein sequences (ie proteins without mutations) are retrieved from UniProt^24^. Amino acid substitutions associated with changes in PPIs are obtained from InAct^25^. We obtained predicted interaction probabilities for canonical and mutants from PLM-interact trained on human data (see Methods). For mutations associated with interaction gain ‘mutation-causing’ interactions (see **Figure 4a**), PLM-interact predicted interaction probabilities of the negative pair are below 0.5, whereas the interaction probabilities of the mutant PPI exceed 0.5. For mutation-disrupting PPIs (**Figure 4b**), the predicted interaction probabilities of the positive PPI exceed 0.5, whereas the interaction probabilities of the mutant PPI are below 0.5. Examples are presented in **Figure 4**, where complex structures are predicted by Chai-1^21^.

**Figure 4.**
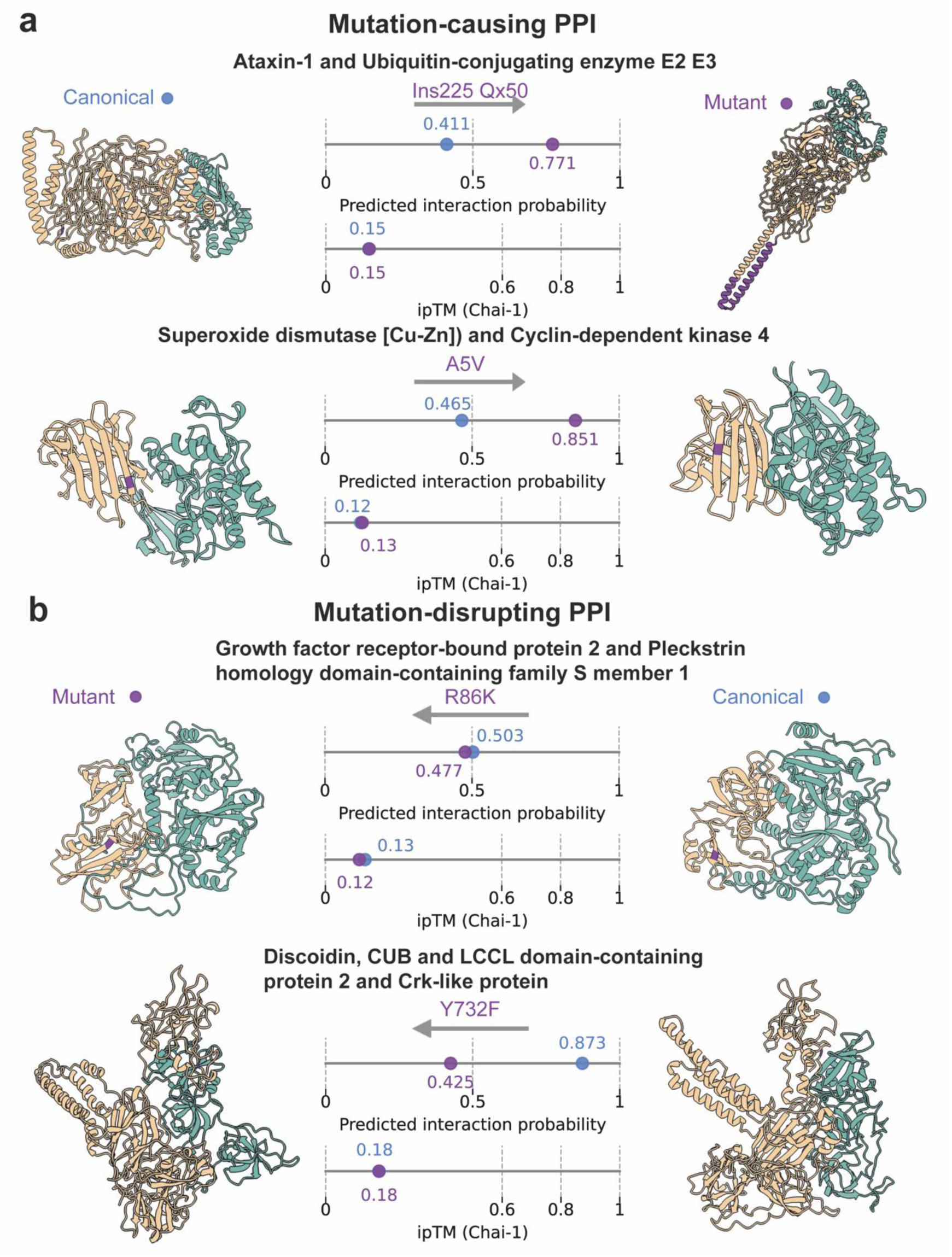
Demonstration of PLM-interact detecting changes in human PPIs associated with mutations. **a** shows two mutation-causing interaction examples, while **b** shows two mutation-disrupting PPI examples. These PPI structures are predicted using Chai-1^21^ and visualised with ChimeraX^23^; here, the mutated amino acids are highlighted in purple. Prediction interaction probabilities exceeding 0.5 indicate the proteins interact, while below 0.5 indicate non-interact. Chai-1’s ipTM scores give the structure prediction confidence where <0.6 indicates failed predictions. Interacting protein structures are shown from left (yellow) to right (green): **a.** Residue 225 Glutamine (Q) of P54253 (Ataxin-1) is mutated to 50 Q, causing interaction with Q969T4 (Ubiquitin-conjugating enzyme E2 E3)^29^; Residue 5 Alanine (A) of P00441 (Superoxide dismutase [Cu-Zn]) is mutated to Valine (V), causing interaction with P11802 (Cyclin-dependent kinase 4)^30^. **b.** Residue 86 Arginine(R) of P62993 (Growth factor receptor-bound protein 2) is mutated to Lysine (K), disrupting its interaction with Q5SXH7-1 (Pleckstrin homology domain-containing family S member 1)^31^; Residue 732 Tyrosine (Y) of Q96PD2 (Discoidin, CUB and LCCL domain-containing protein 2) is mutated to Phenylalanine (F), disrupting its interaction with P46109 (Crk-like protein)^32^. See **Supplementary Figure 4** for AlphaFold3^22^ predicted structures.

We show two examples of mutations that cause PPIs in **Figure 4a**, the protein Ataxin-1 and SOD1 (Superoxide dismutase [Cu-Zn]), which in humans are encoded by the ATXN1 and SOD1 genes separately. Ataxin-1 interacts with many other proteins and the expansion of a glutamine(Q)-encoding repeat can affect the function of PPIs and cause the genetic disease spinocerebellar ataxia type1 (SCA1) and other polyglutamine diseases^26^. Predictions from PLM-interact show that prior to mutation, this PPI has a predicted interaction probability of 0.411, correctly indicating non-interaction with E2s (Ubiquitin-conjugating enzyme E2 E3). Following mutation, PLM-interact increases this score to 0.771, correctly predicting that the mutation induces interaction. In the second example (**Figure 4a**), the SOD1 gene encodes superoxide dismutase enzymes that break down human superoxide radicals. SOD1 is linked to the nervous system disease amyotrophic lateral sclerosis (ALS)^27^, with the A4V mutation being the most common variant in North America^28^. PLM-interact predicts 0.465 and 0.851 before and after the mutation, correctly capturing the change in interaction with CDK4 (Cyclin-dependent kinase 4).

Next, we show two examples of mutations that disrupt PPIs (**Figure 4b**). First, GRB2 (growth factor receptor bound protein 2) is associated with signal transduction. GRB2 mutations are associated with multiple cancers, including breast cancer^33^ and leukaemia^34^. The canonical protein interacts with PLEKHS1 (Pleckstrin homology domain-containing family S member 1). PLM-interact predicts that the missense mutation R86K reduces the interaction probability from 0.503 to 0.477. Second, CLCP1 is a transmembrane protein that regulates cell growth and this protein is identified as a cancer marker, exhibiting up-regulated expression in lung cancer^35^. Again, PLM-interact predicts that the missense mutation Y732F reduces the interaction probability from 0.873 to 0.425.

### Improved virus-human PPI prediction

To study virus-host PPI prediction, we train PLM-interact on a virus-human PPIs dataset from Tsukiyama et al.^11^. The dataset is derived from the Host-Pathogen Interaction Database (HPIDB) 3.0^36^, comprising a total of 22,383 PPIs, which include 5,882 human and 996 virus proteins. We compare our model with three recent virus-human PPI models: PLM-based approach STEP^14^ and the protein embeddings-based approaches LSTM-PHV^11^ and InterSPPI^37^. STEP is similar to existing PPI models benchmarked previously in our study; it leverages protein sequence embedding extracted by the pre-trained PLM ProtBERT^38^. The results show that PLM-interact outperforms the other models. For the STEP comparison this corresponds to improvements in AUPR, F1 and MCC scores of 5.7%, 10.9% and 11.9%, respectively **(Figure 5a**). The length of virus, human proteins and the combined length of virus-human PPIs are shown in **Figure 5b**. To further analyse our model’s performance, we select three pairs of virus-human PPIs from our test data, all with corresponding experimental virus-human complex structures available in the HVIDB^39^. We then use ChimeraX^23^ to visualise these structures and present PLM-interact’s predicted interaction probability for each example (see **Figure 5c**).

**Figure 5.**
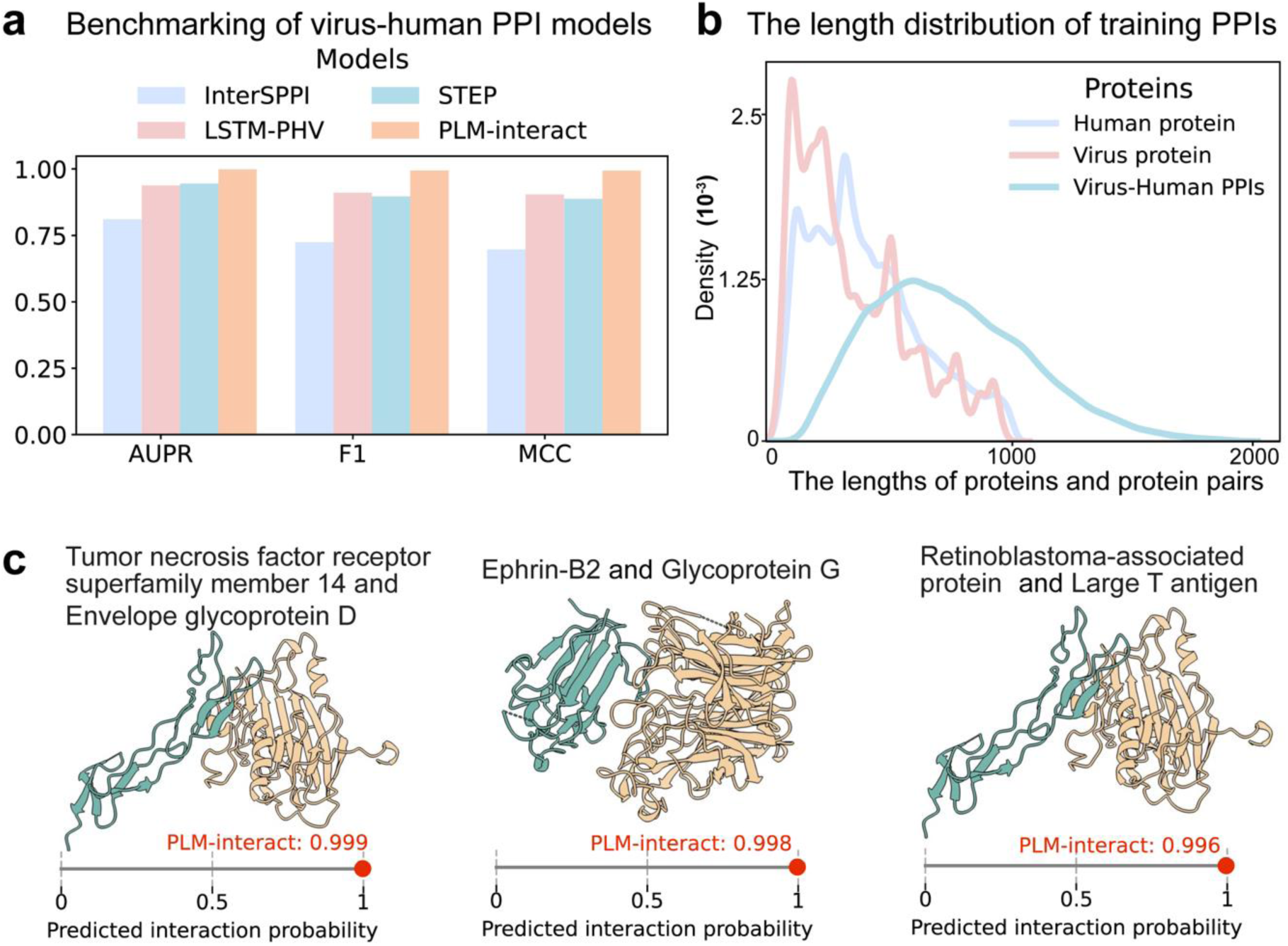
**a.** Comparison of AUPR, F1 and MCC metrics of PLM-interact against recent virus-human PPI models. **b**. The distribution of the length of virus proteins, human proteins and virus-human protein pairs. **c**. The virus-human PPIs are correctly predicted by our model and the 3D complex structures of virus-human PPIs are experimentally verified structures collected from the human-virus PPI database (**HVIDB**^39^). From left (green) to right (yellow), these interacting protein structures are: Tumour necrosis factor receptor superfamily member 14 (Human protein: Q92956) with Envelope glycoprotein D (human herpes simplex virus 1: P57083), Ephrin-B2 (human protein: P52799) with Glycoprotein G (Nipah virus protein: Q9IH62) and Retinoblastoma-associated protein (human protein: P06400) with Large T antigen (Simian virus 40: P03070). Note: The metrics results of the other three models in panel **a** are taken from STEP^14^ paper.

## Discussion

In this study we have developed a novel PPI prediction method, PLM-interact, that extends single protein-focused PLMs to their interacting protein partner. We report significant improvements in held-out species comparisons and highlight successful examples of predicting mutational effects on protein interactions. We further demonstrate PLM-interact’s performance in a virus-human PPI prediction task, showing a significant improvement over state-of-the-art prediction approaches.

Underlying the benefit of PLM-interact is the improved capability of correctly predicting positive PPIs in the held-out species. However, positive PPIs are challenging to predict due to a lack of high-quality PPI data for training. Our improved performance relative to TT3D is particularly impressive given that we only use sequences. TT3D^13^ includes explicit structural information, the per-residual structural alphabet in Foldseek^40^, to improve its prediction over Topsy-Turvy, which incorporated network data.

Furthermore, our results show the potential of predicting mutational effects on PPIs from sequence alone. This could lead to a new generation of interaction aware in-silico variant effect predictors where methods rely on PLMs of the single proteins^41–43^. However, current training data remains limited. The number of high-quality structures of mutant proteins and their interaction partners are low. Algorithmically, models with long and multimodal context^44–46^ that include multiple proteins, structures and nucleotides could be specialised for interaction tasks.

Finally, effective sequence-based virus-host PPI predictors could provide the much-needed molecular detail to conventional virus-host species prediction tools, which tend to rely on genome composition signals ignoring host molecules that are interacting physically with viral molecules^47–50^. In those approaches, the host species only act as labels. Recent progress within SARS-CoV-2 PPI studies mapped out a complex interaction landscape between the virus and human proteome^51,52^. Other viruses are likely to have similarly complex interactions with human and animal hosts. Leveraging these interactions could lead to tools that are better at predicting zoonotic events and the potential for emergence of novel viruses. While PLM-interact has demonstrated significant improvements there is much to do in terms of generating reliable predictions, in particular the need for high-quality virus-host experimental PPI data for training. What is clear is attention applied to longer-range sequence interactions is enhancing our understanding of protein interactions, the fundamental ‘language’ of molecules.

## Methods

### Datasets

The benchmarking human PPI dataset, from Sledzieski et al.^15^, comprises human training and validation data and test data from five other species: mouse, *Mus musculus*; fly, *Drosophila melanogaster*; worm, *Caenorhabditis elegans*; yeast, *Saccharomyces cerevisia*e; and *E. coli*, *Escherichia coli*), all retrieved from STRING V11^53^. We train and validate our model on human PPIs and then conduct inference on PPIs from five other species. All training, validation and test datasets maintain a 1:10 ratio of positive to negative pairs, reflecting the fact that positive PPIs are significantly fewer than negative pairs in the actual host PPI networks. The length of protein sequences ranges from 50 to 800 and PPIs are clustered at 40% identity using CD-HIT^54^ to remove the redundant PPIs. The human training dataset includes 38,344 positive PPIs, whereas the validation set includes 4,794 positive PPIs. Each of the five species includes 5,000 positive interactions, except for *E*. *coli*, which only has 2,000 positive interactions due to the fewer positive PPIs in the STRING dataset used^15^.

The benchmarking dataset of 22,383 virus-human PPIs includes 5,882 human and 996 virus proteins. This dataset was obtained from Tsukiyama et al.^11^, sourced from the HPIDB 3.0 database^36^; the ratio of positive to negative pairs is 1:10 and negative pairs are chosen based on sequence dissimilarities. The length of protein sequences ranges from 30 to 1,000 and the redundant PPIs are filtered based on a threshold of 95% identity using CD-HIT^54^.

In addition, we provide models trained on human PPIs from STRING V12^55^. The positive PPIs are selected by collecting physical links with positive experimental scores, while excluding PPIs with positive homology scores and confidence scores below 400. Previous studies have typically limited the maximum length of protein sequences to 800 or 1000 due to GPU memory limitations. We process the length of protein sequences up to 2000, with a combined length threshold for PPIs of nearly 2500. This human dataset includes 60,308 positive PPIs for training and 15,124 positive PPIs for testing. Furthermore, protein sequences are clustered at 40% identity using MMSeq2^56^ and only PPIs from the distinct clusters are chosen to eliminate redundant PPIs. Again, the positive-to-negative protein pair ratio is 1:10, consistent with the aforementioned two benchmarking datasets.

### Model architecture

We use ESM-2 as the base model in PLM-interact. ESM-2 is an encoder transformer model with a parameter size range from 8 million to 15 billion. The results presented are PLM-interact based on ESM-2 with 650M parameters. We also provided PLM-interact models with ESM-2 35M on our GitHub repository to help with quick testing. The input representation contains amino acid token representations from two proteins. This setup is similar to the original BERT model^57^, also known as the *cross-encoder* which simultaneously encodes a pair of query and answer sentences.

A standard input sequence of PLM-interact, *x*, can be shown as the following:

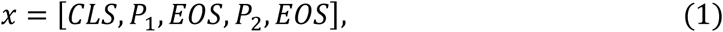

where *CLS* is the classification token, *P*_1_ contains amino acid tokens of protein 1, *P*_2_ contains amino acid tokens of protein 2, and *EOS* is the end-of-sentence token. The first EOS token marks the end of the amino acid sequence in protein 1. This setting allows us to use the original ESM-2 tokenizer to generate embedding vectors *e*, and pass them to the transformer encoder of the ESM-2:

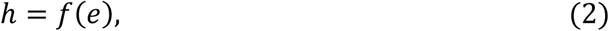

where *f* is ESM-2, *e* contains the token embeddings of *x* , and *h* contains the output embeddings of all input tokens. *h* can be presented as:

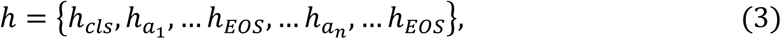

where *h*_*a*1_ and *h*_*an*_ represent amino acid tokens in proteins 1 and 2. Then, we use the *CLS* token embedding to aggregate the representation of the entire sequence pair and as the features for a linear classification function *φ*, and parameterised as a single feed forward layer with a ReLU activation function. The output of the FF layer is converted by the sigmoid function *σ* to obtain the predicted interaction probability *g*,

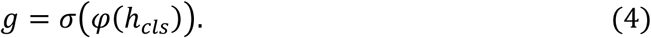

### Model Training

PLM-interact is trained with two tasks: 1) a mask language modelling (MLM) task predicting randomly masked amino acids and 2) a binary classification task predicting the interaction label of a pair of proteins. PLM-interact is trained for 10 epochs using a batch size of 128 on both benchmarking datasets of human PPIs and virus-human PPIs. For all training runs, the input protein pairs are trained on both orders as the interaction between protein 1 and protein 2 is the same as the protein 2 and protein 1, which leads to double the size of the training set. The validation and testing sets are not subject to the same data argumentation. The learning rate is 2e-5, weight decay is 0.01, warm up is 2000 steps, and the schedular is WarmupLinear which linearly increasing the learning rate over the warmup steps. During training, we evaluate the model’s performance at every 2000 steps on the validation set. For every evaluation, a set of 128 protein pairs are randomly sampled from the validation set and the results are averaged over 100 times to ensure metric reliability. Here, we use both masking and classification losses to optimize our model, the loss function for each data point *l* can be represented as:

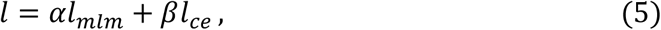

where *l*_mlm_ and *l*_*ce*_ are separately represent the MLM loss and classification (ie cross entropy) loss. *l* can be written as:

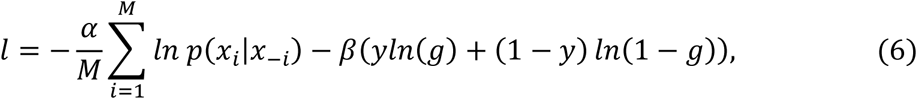

where *M* is the number of the masked tokens, *x*_*i*_ is the true token at position *i* , *p*(*x*_*i*_|*x*_−*i*_) is the probability of the true token *x*_*i*_ given the unmasked amino acid *x*_−*i*_. *y* is the label of the interaction and *g* is the predicted probability for *y* = 1, obtained from equation 4. *⍺* and *β* are weights for the MLM and classification losses.

All of the models are trained on the DiRAC Extreme Scaling GPU cluster Tursa. A typical 10-epoch training run of the model with ESM-2 (650M) with human PPIs takes 31.1 hours on 16 A100-80 GPUs. A typical 10-epoch training run of the model with ESM-2 (650M) trained on virus-human PPIs takes 30.5 hours on 8 A100-80 GPUs. The model with ESM-2 (650M) trained on STRING V12 human PPIs used 16 A100-80 GPUs for 86.4 hours. For model training time with different ratios and model sizes, see the following section optimisation experiment and **Supplementary Table 1**.

We provide model checkpoints that include human PPI models trained on Sledzieski et al.’s benchmarking datasets retrieved from STRING V11^53^, the virus-human PPIs model trained on the benchmarking virus-human PPIs created by Tsukiyama et al.^11^, sourced from HPIDB 3.0^36^, as well as human PPIs collected by us from STRING V12^53^.

### PLM-interact optimisation experiments

To find the optimal value of *⍺* and *β* in equation (6), we benchmark a range of different options between mask loss and classification loss on human benchmarking data. For each ESM-2-35M and ESM-2-650M model, we train five models with different setting of ratios *⍺* : *β* between mask loss and classification loss. The ratios are *⍺* : *β* = 1:1, 1:5, 1:10, 0:1 (with mask), and 0:1 (without mask, denoted as classification). (**Supplementary Figure 1b, c)**. We used the human validation set for each model to identify the optimal epoch checkpoint achieving the best AUPR. Next, the final model is selected based on testing on five other host PPIs.

For ESM-2-650M, the 1:10 ratio is the optimal choice (McNemar test, the greatest counts of p-values less than 0.05) compared to other options (**Supplementary Figure 1d**). The model with a 1:5 ratio of mask to classification loss based on ESM-2-35M achieved the highest number of significant improvements (McNemar test, the greatest counts of p-values less than 0.05) compared to other models (**Supplementary Figure 1e**). According to these results, we select a loss ratio of 1:10 for ESM-2-650M and 1:5 for ESM-2-35M. The ratio setting is implemented in both benchmarking of human PPI training and virus-human PPI training, as well as human PPI training using the STRING V12 database.

### Baselines

We compute the prediction interaction probabilities based on checkpoints of TT3D^13^, Topsy-Turvy^19^ and D-SCRIPT^15^ to generate precision-recall (PR) curves in **Figure 2b**. Due to the absence of publicly available checkpoints for DeepPPI^20^ and PIPR^6^, these methods are excluded from the PR curve comparison. The AUPR values for DeepPPI and Topsy-Turvy are sourced from the Topsy-Turvy paper^19^, those for D-SCRIPT and PIPR are from the D-SCRIPT paper^15^, and the AUPR value of TT3D is obtained through email communication. A complete list of the main features, architectures, reference, and code link of each baseline method can be found in **Supplementary Table 2**.

### MMseq2

We used MMseq2 to obtain the protein sequence-based alignment results between each pair of proteins; the parameters setting is:

--threads 128 --min-seq-id 0.4 --alignment-mode 3 --cov-mode 1.

### Chai-1

Chai-1^21^ is a state-of-the-art model for molecular structure prediction, available at https://lab.chaidiscovery.com/. We use Chai-1 with the “specify restraints” option to predict protein-protein structure complexes and visualise predicted PPI structures using the molecular visualisation program ChimeraX^23^.

### AlphaFold3

AlphaFold3 is a tool to predict the biomolecular interactions including protein, DNA, small molecules, ions and modified residues^22^, available at https://alphafoldserver.com/. We use AlphaFold3 in its PPI mode to predict protein structure complexes. The results are visualised with the molecular visualisation program ChimeraX^23^.

### McNemar’s test

McNemar’s Chi-Square test^58^ is a statistical test that determines if there are significant differences between paired nominal data.

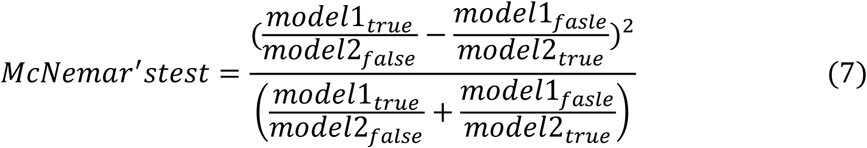

Here, *model*1_*true*_and *model*2_*true*_separately represent the count of correct prediction obtained by *model*1 or *model*2, while *model*1_*fasle*_and *model*2_*false*_separately represent the count of incorrect prediction obtained by *model*1 and *model*2.

To investigate if our models with varying ratios of mask-to-classification loss perform significant differences, we conducted a McNemar’s test for any pairs of models in optimisation experiments. This test is based on the number of correct and incorrect between two models. Predicted interaction probabilities from each model are used to get predicted labels, which are used to obtain the counts of correct and incorrect predictions. A McNemar’s test p-value ≤ 0.05 indicates a significant difference between the predictive performance of two models. The model with more correct predictions is considered superior to the other.

## Data availability

Sledzieski et al.’s benchmarking PPI data is available at https://d-script.readthedocs.io/en/stable/data.html

Tsukiyama et al.’s virus-human benchmarking PPIs dataset is available at: http://kurata35.bio.kyutech.ac.jp/LSTM-PHV/download_page

STRING V12 PPIs database: https://stringdb-downloads.org/download/protein.physical.links.v12.0.txt.gz

The mutations that cause or disrupt PPIs are from IntAct^25^ Database; the link is https://ftp.ebi.ac.uk/pub/databases/intact/current/various/mutations.tsv

UniProt: https://www.uniprot.org/

HVIDB: http://zzdlab.com/hvidb/download.php

## Code availability

The code and trained models are available at https://github.com/liudan111/PLM-interact

## Acknowledgements

The authors acknowledge funding from the European Union’s Horizon 2020 research and innovation 562 program, under the Marie Sklodowska-Curie Actions Innovative Training Networks 563 grant agreement no. 955974 (VIROINF) for DL, a UK Medical Research Council (MRC) Doctoral Training Programme in Precision Medicine studentship (MR/N013166/1) for KDL and MRC grants: MC_UU_00034/5, MC_UU_00034/6 and MR/V01157X/1. KY acknowledges support from Cancer Research UK (EDDPGM-Nov21\100001 and DRCMDP-Nov23/100010), Biotechnology and Biological Sciences Research Council (BBSRC) BB/V016067/1, Prostate Cancer UK MA-TIA22-001 and EU Horizon 2020 grant ID 101016851. This work used the DiRAC Extreme Scaling service (Tursa) at the University of Edinburgh, managed by the Edinburgh Parallel Computing Centre on behalf of the STFC DiRAC HPC Facility (www.dirac.ac.uk). The DiRAC service at Edinburgh was funded by BEIS, UKRI and STFC capital funding and STFC operations grants. DiRAC is part of the UKRI Digital Research Infrastructure.

## Supplementary Figures

**Supplementary Figure 1.**
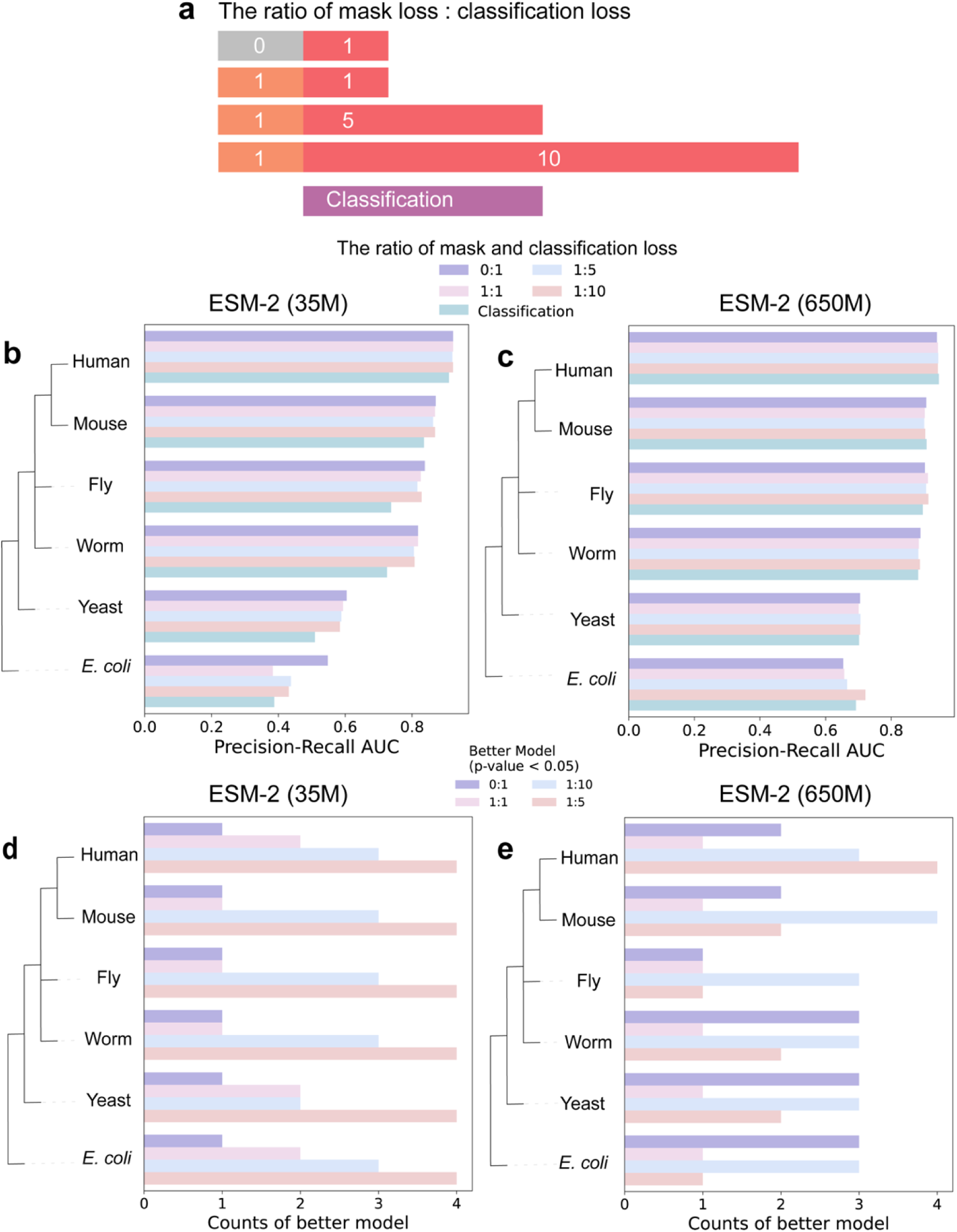
The benchmarking of different ratios of mask to classification loss on five species PPI prediction. **a.** The bar plot to show the ratio between mask loss and classification loss. **b** and **c** respectively represent the performance of our model with the different ratios between mask and classification loss on 650M and 35M of ESM-2 models. The left is aligned with taxonomy tree of the hosts that are used for evaluating our human PPI model. **d** and **e** show the better model with significant improvements than the other, p-value < 0.05 with McNemar’s Test.

**Supplementary Figure 2.**
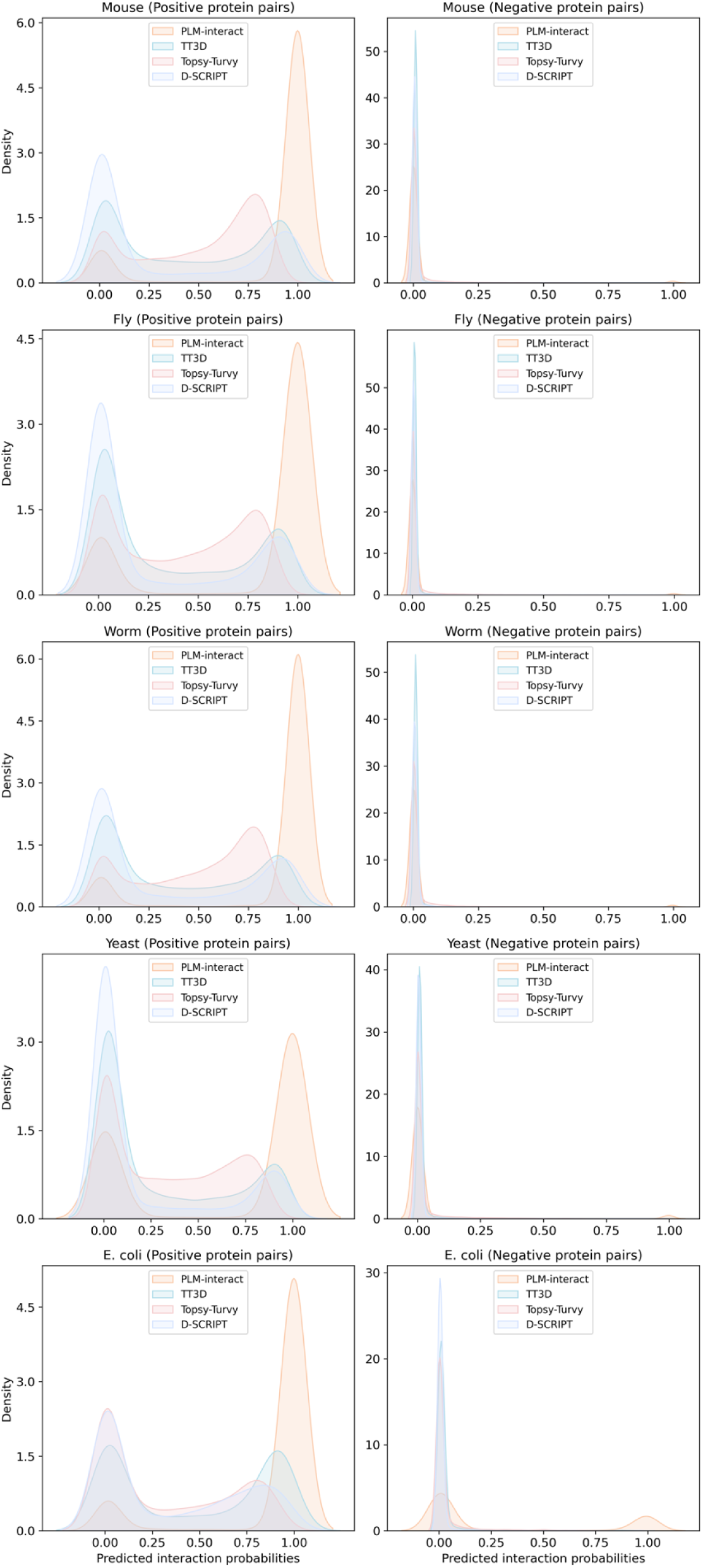
The distribution of prediction scores of positive and negative protein pairs of PLM-interact, TT3D, TT and D-SCRIPT. PLM-interact outperforms other models by identifying the most true positive pairs (predicted interaction probability > 0.5) and true negative pairs (predicted interaction probability < 0.5), demonstrating that PLM-interact achieves the best performance, except for negative pairs of *E. coli*.

**Supplementary Figure 3.**
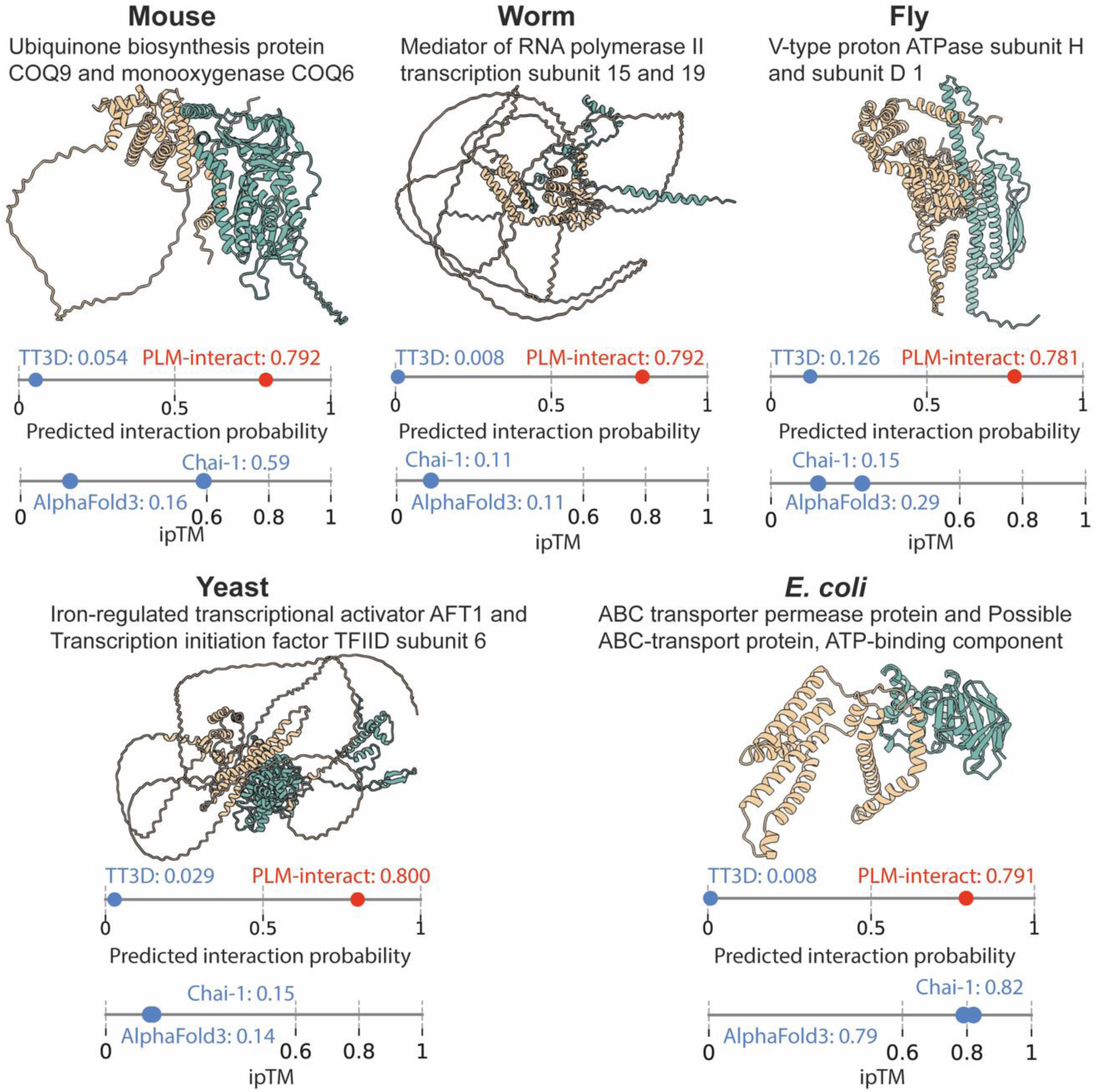
PPI example for each species that was predicted correctly by PLM-interact but not by TT3D. Protein-protein structures are predicted by AlphaFold3^22^. and visualised with ChimeraX^23^. Both models’ prediction interaction probabilities range between 0 and 1. A predicted interaction probability >0.5, is predicted as a positive PPI, while <0.5 is a negative pair. Interacting proteins are shown from left (yellow) to right (green), respectively. For information about these PPIs, see Figure 3.

**Supplementary Figure 4.**
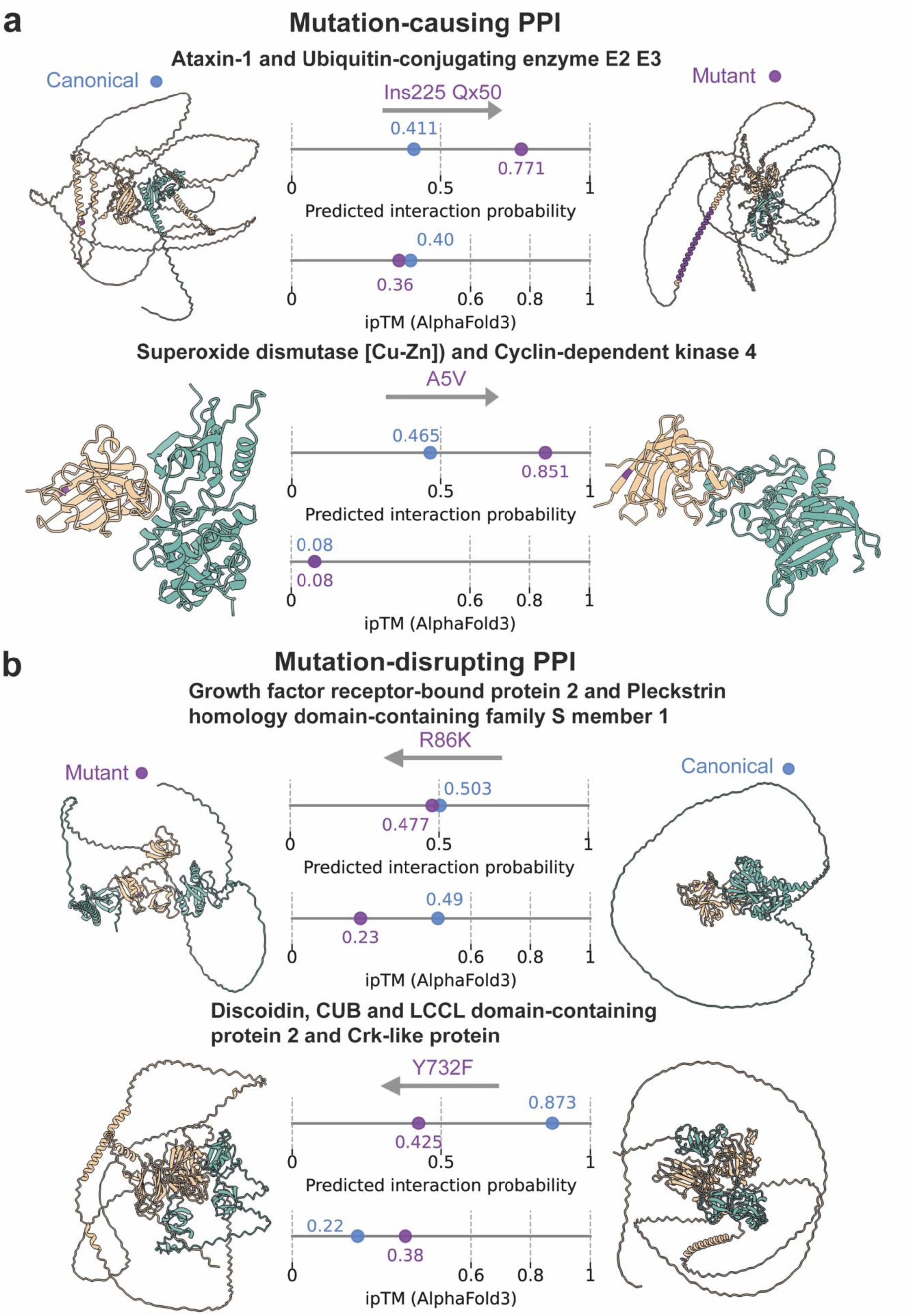
Demonstration of PLM-interact detecting changes in human PPIs associated with mutations. **a** shows two mutation-causing interaction examples, while **b** shows two mutation-disrupting PPI examples. These PPI structures are predicted using AlphaFold3^22^ and visualised with ChimeraX^23^; here, the mutated amino acids are highlighted in purple. Prediction interaction probabilities exceeding 0.5 indicate the proteins interact, while below 0.5 indicate non-interact. AlphaFold3’s ipTM scores give the structure prediction confidence where <0.6 indicates failed predictions. Interacting protein structures are shown from left (yellow) to right (green). See Figure 4 for information about these protein pairs.

## Supplementary Tables

**Supplementary Table 1.** This table shows the GPU hours (GPUhs) of different models. PLM-interact has the two kinds of training strategies: masking language modelling and binary classification, binary classification only. The ratios in this table present the different weights between mask loss and classification loss, binary indicates the binary classification task.

| Models/GPUhs | 650M | 35M |
| --- | --- | --- |
| 0:1 | 522.4 | 238.2 |
| 1:1 | 514.3 | 299.1 |
| 1:5 | 508.4 | 238.1 |
| 1:10 | 496.9 | 240.4 |
| Binary | 355.8 | 130.9 |

**Supplementary Table 2.**
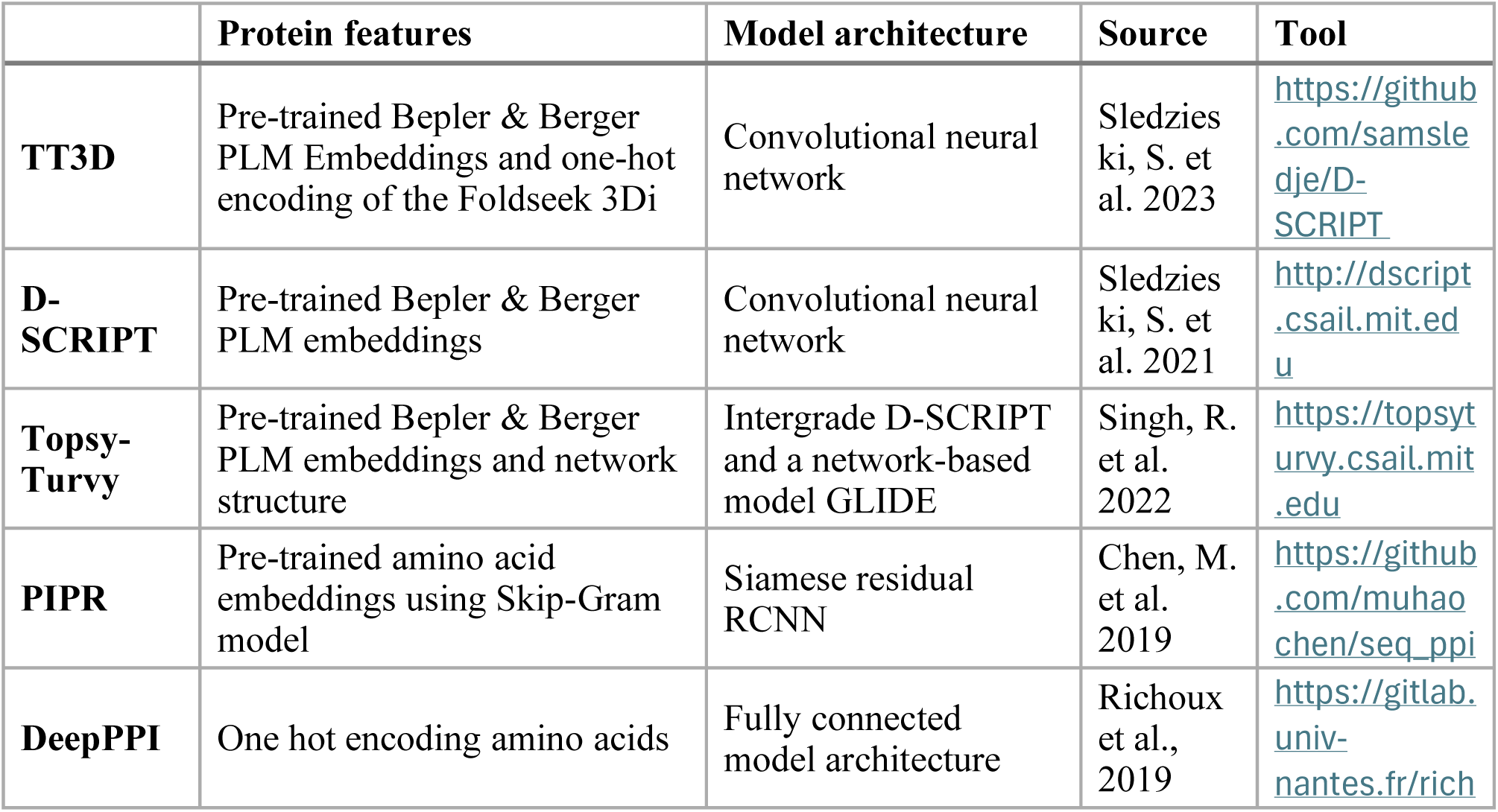
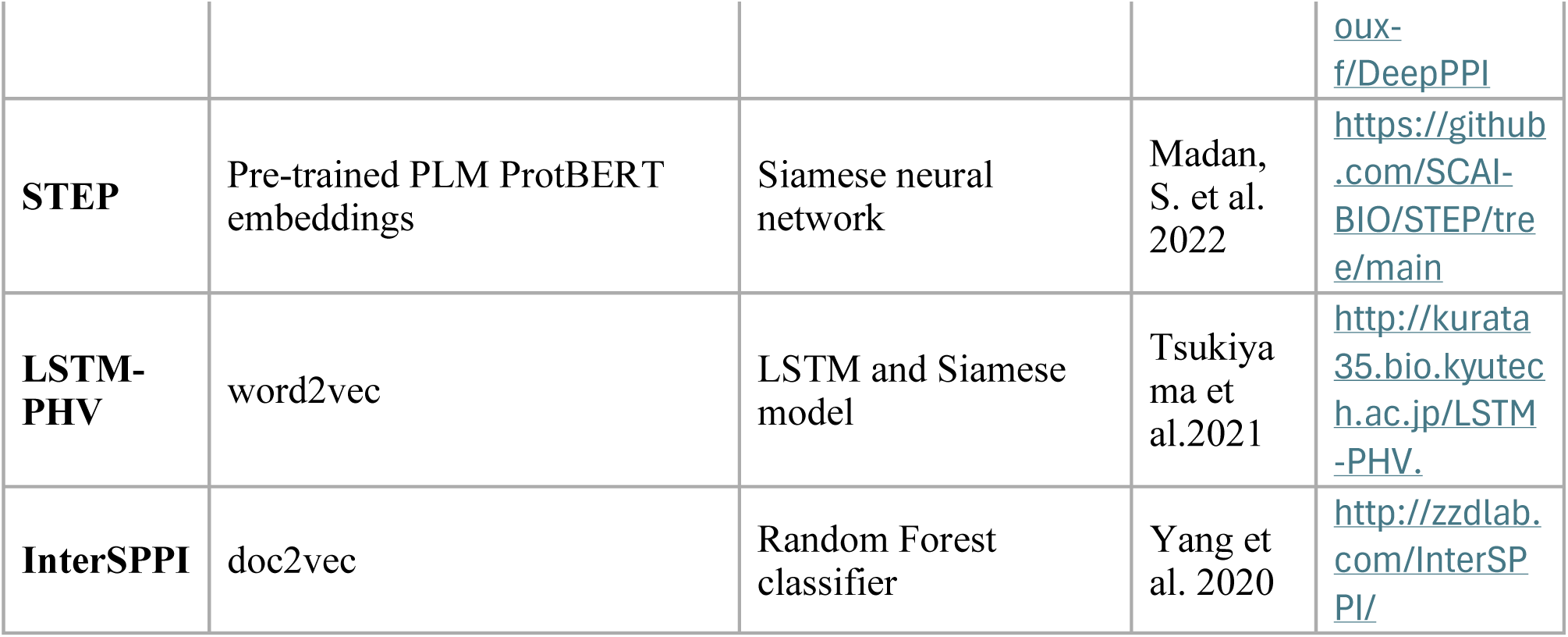
This table described the features of state-of-the-art models in this paper.

